# Novel role for peptidoglycan carboxypeptidases in maintaining the balance between bacterial cell wall synthesis and degradation

**DOI:** 10.1101/2023.07.12.548665

**Authors:** Manuela Alvarado Obando, Tobias Dörr

## Abstract

Peptidoglycan (PG) is the main component of the bacterial cell wall; it maintains cell shape while protecting the cell from internal osmotic pressure and external environmental challenges. PG synthesis is essential for bacterial growth and survival, and a series of PG modifications are required to allow expansion of the sacculus. Endopeptidases (EPs), for example, cleave the crosslinks between adjacent PG strands to allow the incorporation of newly synthesized PG. EPs are collectively essential for bacterial growth and must likely be carefully regulated to prevent sacculus degradation and cell death. However, EP regulation mechanisms are poorly understood. Here, we used TnSeq to uncover novel EP regulation factors in *Vibrio cholerae*. This screen revealed that the carboxypeptidase DacA1 (PBP5) alleviates EP toxicity. *dacA1* is essential for viability on LB medium, and this essentiality was suppressed by EP overexpression, revealing that EP toxicity both mitigates, and is mitigated by, a defect in *dacA1*. A subsequent suppressor screen to restore viability of *ΔdacA1* in LB medium was answered by hypomorphic mutants in the PG synthesis pathway, as well as mutations that promote PG degradation. Our data thus reveal a key role of DacA1 in maintaining the balance between PG synthesis and degradation.

## Introduction

The bacterial cell wall serves as a vital structure for bacterial growth and survival. It fulfils multiple functions, including acting as a protective layer that imparts structural stability, facilitating adaptation to drastic environmental changes, and determining cell shape. The cell wall is composed of peptidoglycan (PG), a mesh-like structure made up of alternating subunits of N-acetyl glucosamine (GlcNAc) and N-acetylmuramic acid (MurNAc), with each MurNAc unit bearing a short peptide stem composed of 5 amino acids (L-ala, D-glu, mDAP, D-ala, D-ala). PG expansion during growth is mediated by two key synthetic processes, transglycosylation (TG) and transpeptidation (TP) reactions. TG and TP reactions are performed by PG synthases like penicillin-binding-proteins (PBPs) and shape, elongation, division and sporulation (SEDS) proteins^1^. During transglycosylation, the PG strands are polymerized by forming β-(1,4) glycosidic bonds. During or after PG strand polymerization, the transpeptidation reaction is used to form crosslinks between adjacent chains via the formation of peptide bonds between most commonly the mDAP residue from one peptide stem (the “acceptor”) and the 4^th^ D-ala of another (the “donor”); this results in loss of the terminal (5^th^) D-ala from the donor strand (while, importantly, the acceptor strand can retain the 5^th^ D-ala).

Besides PG synthesis, multiple tailoring modifications are required to produce a mature and flexible cell wall. These modifications are carried out by a group of enzymes collectively referred to as “autolysins”, which collectively can cleave almost every linkage in the PG structure. Autolysins include amidases, lytic transglycosylases, carboxypeptidases and endopeptidases. Carboxypeptidases remove the terminal D-ala of the pentapeptide chain of newly synthesized peptidoglycan (which is where pentapeptide is expected to occur exclusively). Their proposed function is to control the amount of pentapeptide subunits that are available for crosslinking by transpeptidation, but how exactly this contributes to proper growth and morphogenesis is unclear^2^. In *E.coli* DacA (PBP5) is required for maintaining cell diameter, surface uniformity and overall topology of the peptidoglycan sacculus^3^, however, the mechanisms behind the morphological aberrations that occur when DacA1 is absent have not been elucidated.^4^ In *V. cholerae*, DacA1 deletion results in drastically impaired growth and morphology in NaCl-containing medium (LB) and DacA1 is consequently (for an unknown reason) essential for viability in high-salt, but not low-salt conditions ^5^.

Another prominent autolysin group, the endopeptidases, cleave the crosslinks between peptide side stems of two adjacent PG strands. This activity is thought to be required for the directional insertion of new PG material during cell elongation (and probably division); consequently, EP activity is essential for growth and viability in all bacteria studied in this regard to date ^6–8^ Interestingly, EPs represent a double-edged sword for the cell, as they are at once essential for cell growth, but also potentially detrimental to the structural integrity of the PG sacculus if not properly regulated. Tight regulation of endopeptidase activity at an optimal level is therefore essential to maintain cell growth and morphology. Endopeptidase regulation mechanisms have only begun to be understood in gram-negative bacteria; proposed mechanisms include regulation at the transcriptional level^9^, proteolytic degradation^10^, substrate selectivity^6^ and post-translational activation through conformational changes^11^. The latter mechanism is an important aspect of endopeptidase regulation in *V. cholerae’s* main endopeptidase ShyA, where conformational re-arrangements allow ShyA to transition from a closed (inactive) conformation where domain 1 occludes the catalytic groove of domain 3, to an open (active) conformation where this inhibition is relieved. The factors involved in this conformational switching remain unknown. Mutating residues in the domain 1/domain 3 interaction surface (most prominently the L109K change) causes disruption of hydrophobic interactions between domain 1 and 3, resulting in a more open protein conformation and subsequent hyperactivity. How EP activity is harnessed to properly direct PG insertion during cell elongation, and how EPs intersect with other autolysins, is largely unknown.

Here, we reveal a new functional interaction between carboxypeptidases and EPs. We employed TnSeq to identify factors that exacerbate or mitigate toxicity of an activated EP and discovered the carboxypeptidase DacA1(PBP5) as a previously unknown alleviator of cell wall degradation. Surprisingly, *dacA1* itself is essential for growth and this was suppressed by EP overexpression. A screen for additional suppressors of *dacA1* essentiality revealed mutants in the PG synthesis pathway that exhibit decreased function, as well as mutations that promote PG degradation. Our findings thus suggest a key role for DacA1 in preserving the equilibrium between PG synthesis and degradation.

## Results

### A screen for endopeptidase regulators identifies a genetic relationship between ShyA and DacA1 (PBP5)

As part of an ongoing effort to identify regulators of peptidoglycan endopeptidase activity, we conducted a Transposon Sequencing Screen (TnSeq) in *V. cholerae* to identify genes that modulate the toxicity of the activated version of EP ShyA (ShyA^L109K^). Overexpression of ShyA^L109K^ exhibits enhanced cleavage activity due to the mutational relief of an intramolecular inhibitory mechanism^12^. We thus created Tn insertion libraries (3 x 200,000 colonies) in a strain background that overexpresses ShyA^L109K^ from a chromosomal locus and identified genome-wide Tn insertion frequencies vs. WT control. The screen identified several Tn insertion events that were significantly underrepresented in the ShyA^L109K^ strain (indicating conditional essentiality in this background), as well as some that were significantly overrepresented, indicating genes whose inactivation reduces ShyA^L109K^ toxicity.

Among the conditionally essential genes, this screen was answered by PG synthesis and recycling factors such as the main PBP, *pbp1A,* as well as its accessory factor *csiV*^13^ and several PG recycling pathway components, namely the muropeptide ligase *mpl,* the muropeptide fragment permease *ampG*^14^ and the lytic transglycosylase *mltB* (Fig. 1A). Another synthetic lethal hit was *yhdP*, which likely fulfils phospholipid transport functions to the OM, with a preference for saturated fatty acids^15^. ShyA^L109K^ overexpression causes spheroplast formation, and spheroplast survival has been shown to depend on YhdP^16^; this hit may thus suggest that saturated fatty acids contribute to spheroplast stability. As internal validation, insertions in *shyA* answered the screen as synthetic healthy, likely reflecting insertions into the chromosomal ShyA^L109K^ overexpression construct (Fig. 1AB). Intriguingly, the carboxypeptidase DacA1 was conditionally enriched in our TnSeq, indicating a synthetic healthy relationship between *dacA1* and *shyA^L109K^*(Fig. 1B). The deletion of *dacA1* in cells overexpressing ShyA^L109K^ resulted in complete rescue of cell growth, validating the TnSeq results (Fig. 1C). This indicates that DacA1, the main carboxypeptidase in *V. cholerae,* either directly or indirectly promotes toxicity of ShyA^L109K^, suggesting an uncharacterized link between CPase and EP activity. We were intrigued by this genetic relationship and chose to characterize this connection further.

**Figure 1.**
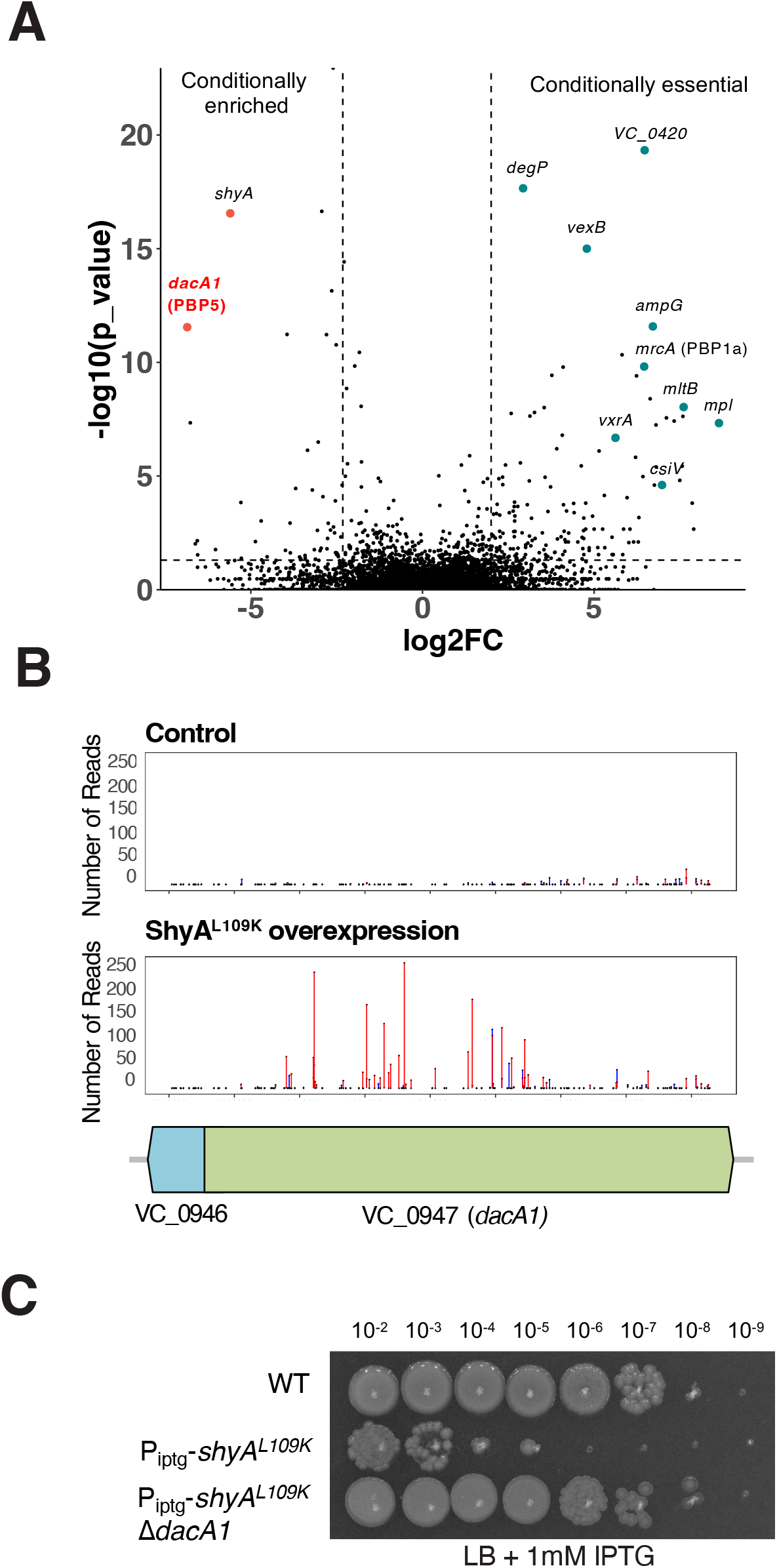
A screen for EP regulators identifies a genetic relationship between ShyA and PBP5. **(A)** Volcano plot of the change in relative abundance of insertion mutants between the control condition (WT) and the experimental condition (ShyA^L109K^ overexpression) (X-axis) versus p-value (Y-axis). The dashed line indicates the cutoff criterion (>2-fold fitness change, P <0.005) for identification of genes that modulate ShyA^L109K^ toxicity. **(B)** *dacA1* was identified as conditionally enriched upon ShyA^L109K^ overexpression using CON-ARTIST, red bars represent the number of reads from transposon insertions found in the forward orientation and blue bars represent insertions found in the reverse orientation. The black dots indicate every TA site available for transposon insertion **(C)** Overnight cultures of WT, P_iptg_-shyA^L109K^ and P_iptg_-shyA^L109K^ *ΔdacA1* where plated on LB medium with 1mM IPTG and incubated overnight at 30°C.

### PG of *ΔdacA1* mutant is less susceptible to ShyA cleavage

We have previously shown that ShyA exhibits substrate selectivity, i.e. it prefers cleaving crosslinks that contain tetrapeptides over those that contain pentapeptides, and speculated that this represents a strategy to differentiate between old and new PG^6^. We have also shown that DacA1 is the main carboxypeptidase under normal laboratory conditions, as its deletion results in a >8-fold increase in pentapeptide content^5^. We thus hypothesized that the increase in pentapeptide content in the absence of DacA1 causes reduced susceptibility to ShyA^L109K^ cleavage, thus conferring resistance against EP toxicity (Fig. 2A). To test this hypothesis, we purified ShyA and ShyA^L109K^, and used a Remazol-blue PG degradation assay to quantify PG hydrolysis. As substrate, we used PG purified from either WT or *ΔdacA1* cells (Fig. 2B). The activity of both ShyA and ShyA^L109K^ was significantly reduced on Δ*dacA1* PG compared to WT PG, suggesting that indeed Δ*dacA1* cell wall is at least partially refractory to EP degradation. To exclude the possibility that Δ*dacA1* PG is somehow generally less susceptible to enzymatic degradation (through, e.g. non-specific PG architectural traits), we also included lysozyme as a control, which should be active regardless of pentapeptide content due to its activity on PG’s polysaccharide backbone. Indeed, lysozyme activity was unaffected by PG origin (Fig 2C). These results indicate that modifications in the PG side stem structure generated by DacA1 (likely pentapeptide) reduce ShyA activity, suggesting a role for DacA1 in substrate-guided endopeptidase regulation.

**Figure 2.**
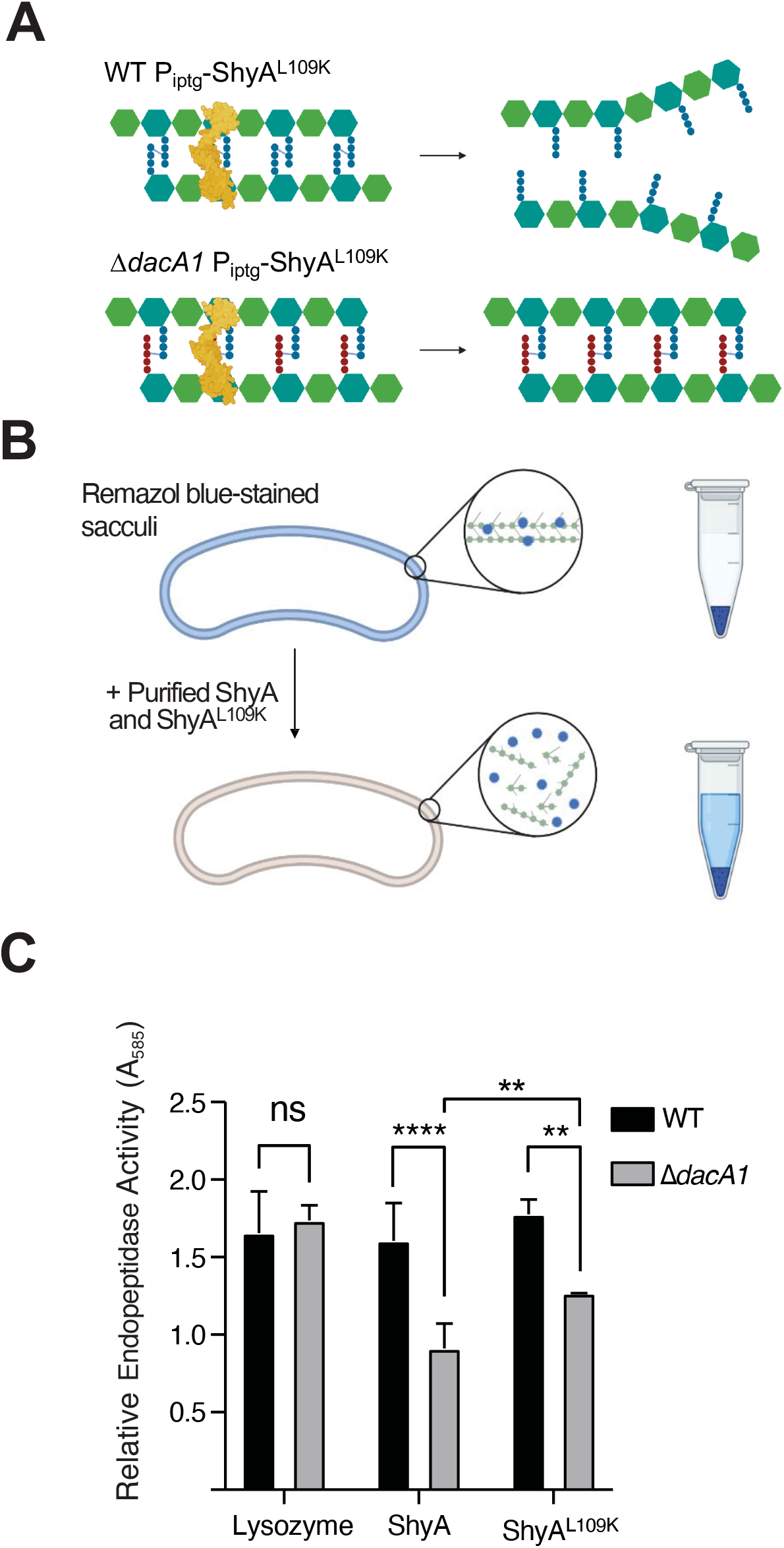
Δ*dacA1* mutant PG is less susceptible to cleavage by ShyA. **(A)** Schematic representation of how DacA1 absence affects the pentapeptide content in the PG sacculus and as a result potential ShyA activity. **(B)** Schematic of experimental procedure. The PG sacculi from WT and *ΔdacA1* cells were purified using the SDS precipitation method^47^. Purified samples were then stained with Remazol Blue and used in sedimentation assays where PG fragmented by hydrolases remains soluble in the supernatant upon centrifugation, providing a quantitative readout of EP activity **(C)** RBB stained sacculi were treated as described in (B) with the addition of proteins indicated on the x-axis. Absorbance values were normalized against the buffer. Error bars represent standard deviation of two independent experiments. Significance was determined using a 2way ANOVA, **** corresponds to a P-value of <0.0001.

### Upregulation of cell wall degradation functions suppresses ΔdacA1 essentiality

The finding that a *dacA1* mutant rescued ShyA^L109K^ overexpression toxicity was unexpected, since DacA1 itself is essential for growth in LB (the growth medium in which we conducted our screen), but non-essential for growth in salt-free LB^5^. The absence of DacA1 in *V. cholerae* results in severe growth and morphological defects when grown in LB medium, and these phenotypes specifically depend on the NaCl concentration, rather than osmolarity^5^. When validating the genetic relationship between ShyA^L109K^ and DacA1, we found that increased endopeptidase activity from ShyA^L109K^ rescued the *ΔdacA1* growth and morphology defects (Fig. 3ABC, Fig. S1). Thus, curiously, the Δ*dacA1* mutation and ShyA^L109K^ overexpression are two individually toxic events that suppress each other’s detrimental effects on the cell. This observation suggests that reduction in PG degradation might contribute to *ΔdacA1* defects in LB. To address the genetic basis of the *ΔdacA1* salt sensitivity phenotype, and thus potentially further characterize its relationship with ShyA, we performed three independent genetic screens to select for suppressor mutations that rescued the Δ*dacA1* mutant’s growth on LB medium and mapped them via whole genome sequencing. A common pattern in the mutations that answered this screen was an enrichment in mutants associated with either cell wall degradation, or synthesis, elaborated in further detail below.

**Figure 3.**
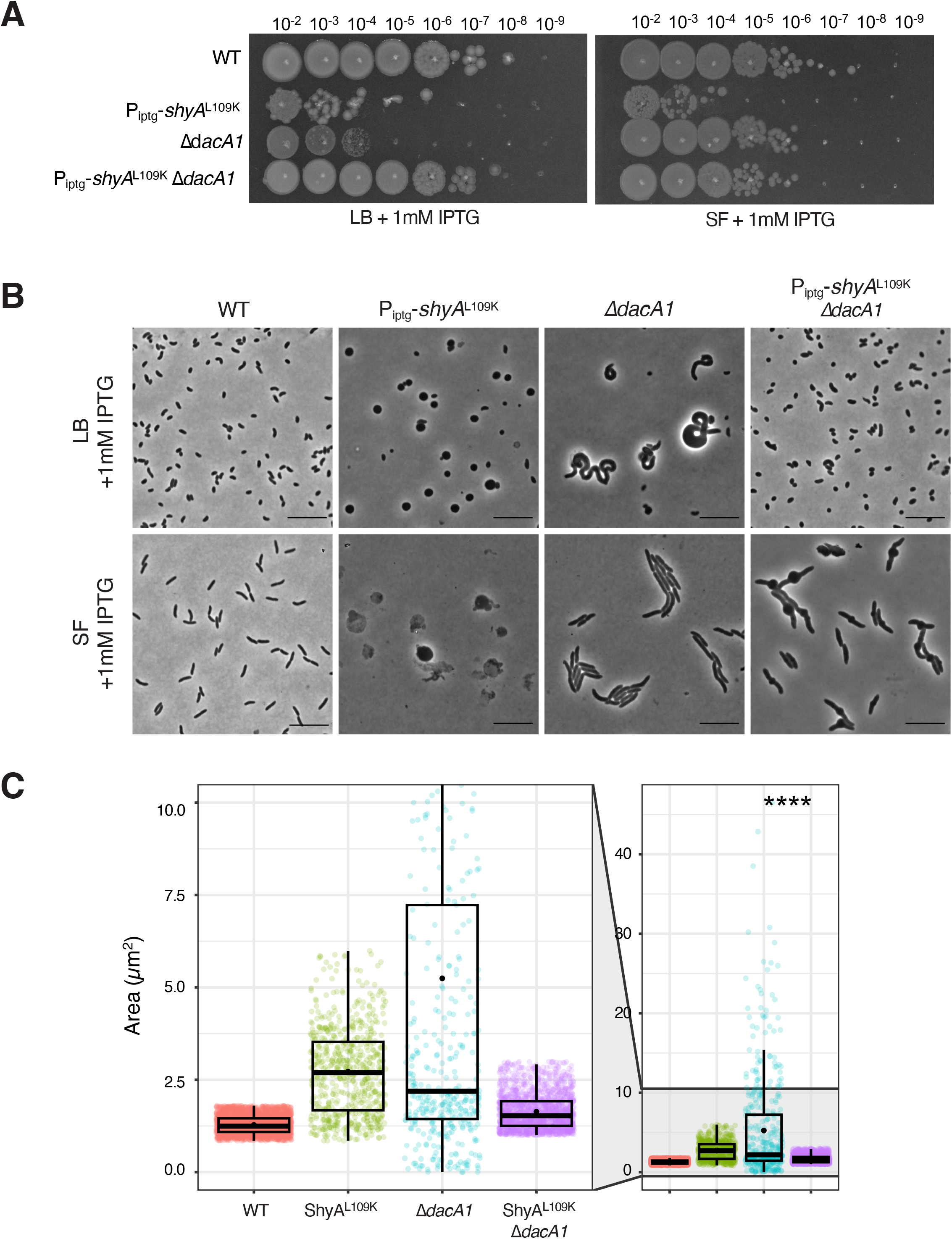
ShyA^L109K^ overexpression restores normal cell growth and morphology in Δ*dacA1* cells grown on LB media. **(A)** Overnight cultures were grown on salt-free LB and then spot-tittered on either LB or salt-free LB with 1mM IPTG. **(B)** Overnight cultures were sub-cultured 1:100 in either LB or salt-free LB at 37°C until they reached exponential phase, followed by addition of inducer (1mM IPTG). Phase contrast images were taken after 2 hours of induction. **(C)** Cells were segmented using Omnipose, and area, width and length were calculated with MicrobeJ. Statistical significance was determined with Welch’s two sample t-test. ****, P < 0.0001. Shown are boxplots with raw data points.

Among the genes that are associated with cell wall degradation, we found suppressor mutations in two genes that are expected to result in endopeptidase activation. One mutation mapped to *zur*, a Fur family transcriptional regulator known to repress the expression of ShyB - another endopeptidase in *V. cholerae* - under zinc replete conditions^9^ (Fig. 4A). We hypothesized that the *zur* mutation represented a loss of function allele, resulting in upregulation of ShyB. To test this, we constructed a clean deletion of *zur* in a *ΔdacA1* background. This resulted in partial rescue of the *ΔdacA1* phenotype in LB (Fig. 4A). We then confirmed that partial rescue was due to *zur* deletion and not a polar effect by expressing *zur in trans*. Next, we constructed a strain where ShyB was under the control of an IPTG inducible promoter to test more directly its ability to suppress Δ*dacA1* growth defects. Indeed, ShyB expression rescued *ΔdacA1* normal growth (Fig. 4B) as well as morphological defects (Fig. 4E) (Fig. S2) in LB medium, showing that upregulation of another EP can suppress Δ*dacA1* essentiality.

**Figure 4.**
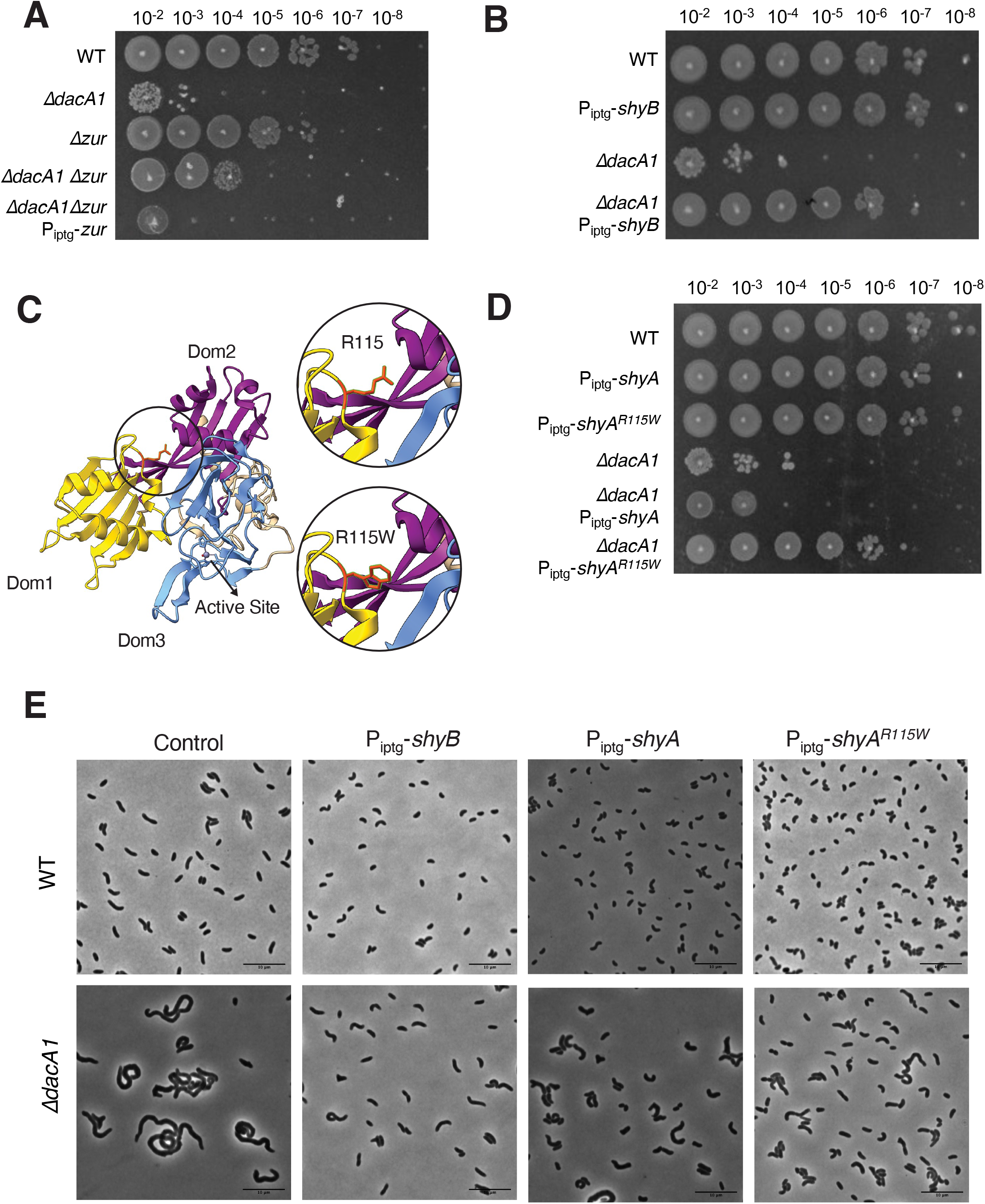
Mutations that enhance endopeptidase activity suppress the *ΔdacA1* phenotype in LB. **(A)** Overnight cultures of the indicated strains were plated by spot-tittering in LB media and incubated at 30°C overnight. **(B)** Overnight cultures of the indicated strains were sub-cultured 1:100 and grown at 37°C in LB medium until they reached exponential phase, followed by addition of inducer (1mM IPTG) for 3 hours. Cells were then plated on LB medium containing 1mM IPTG **C)** Crystal structure of ShyA showing residue R115 in the interphase between Dom 1 and Dom 3, and prediction of the R115W mutation. **(D)** Mutant ShyA^R115W^ was overexpressed in WT and *ΔdacA1* backgrounds, and induction was performed as described in Figure 4B legend. **(E)** Overnight cultures were sub-cultured 1:100 in either LB or salt-free LB at 37°C until they reached exponential phase, followed by addition of inducer (1mM IPTG). Phase contrast images were taken after 3 hours of induction.

Most intriguingly, the screen was also answered by a mutation in ShyA (R115W). Residue R115 lies in the interaction surface between ShyA’s inhibitory domain 1 and its active site domain 3 (Fig. 4C), close to many residues we have previously shown to activate ShyA activity^11^. Consistent with an activating role of this mutation, ShyA^R115W^ exhibited increased toxicity (reduced plating efficiency and morphological defects) when expressed in *E. coli* (Fig. S3). We then overexpressed ShyA^R115W^ from an IPTG inducible promoter in the WT and Δ*dacA1* backgrounds. Unlike ShyA^L109K^, overexpression of ShyA^R115W^ did not affect growth or produce spheroplasts in a *V. cholerae* WT background; however, it was still able to partially rescue growth and morphology in the *ΔdacA1* phenotype in LB medium (Fig. 4D,E)(Fig. S2). Taken together, these results indicate that increased endopeptidase activity is beneficial for the *ΔdacA1* mutant in LB medium, suggesting that the mutant suffers from reduced cell wall turnover as the underlying cause of its growth and morphology defects.

### Downregulation of cell wall synthesis suppresses ΔdacA1 essentiality

The second category of mutants that answered the screen mapped to genes involved in PG precursor synthesis (*murA^P122S^*, *murA^L35F^ murC^A132T^* and *murD^D447E^)*. Since *murA murC* and *murD* are essential for growth, these mutations must represent either hypomorphs or gain-of-function mutations, rather than loss of function. Based on our finding that increased PG hydrolysis rescued the Δ*dacA1* mutant, we hypothesized that these *mur* mutations may accomplish something similar by reducing PG synthesis, i.e. that they are hypomorphs. The mutations in MurA and MurC mapped close to the active site, indeed suggesting they may result in reduced activity, while the *murA* mutations require more complex interpretation. The MurA structure consists of two globular domains, with its active site located in the interface between them^17^. In *E. coli,* residue C115 has been identified as the catalytic cysteine required for MurA activity as is often targeted by MurA inhibitors^18^. Residue P122, which was mutated to serine in one of our suppressors, is situated in the interface between both domains and near the active site. Proline is the most rigid amino acid^19^, therefore replacing it with a serine (P122S) will likely induce a significant conformational change affecting MurA activity. The other mutation (L35F) is located in the N-terminal domain of the protein, and its effect is thus harder to gauge; however, reports in other organisms suggest mutations in the globular domains may have a role in MurA turnover by ClpCP^20^ and protein-protein interactions^21^. (Fig. S4). To test the role of these mutations in suppressing Δ*dacA1* defects further, we first reconstituted the *murA^P122S/L35F^* and *murD^D447E^*mutations in naïve WT and Δ*dacA1* backgrounds (we were unable to construct *murA^P122S^* in the wild-type background). All three mutations restored colony formation of *ΔdacA1* on LB (Fig. 5A); however, further investigation revealed some complexity. For example, neither mutation restored the morphological defects; rather, they exacerbated the division defect.

**Figure 5.**
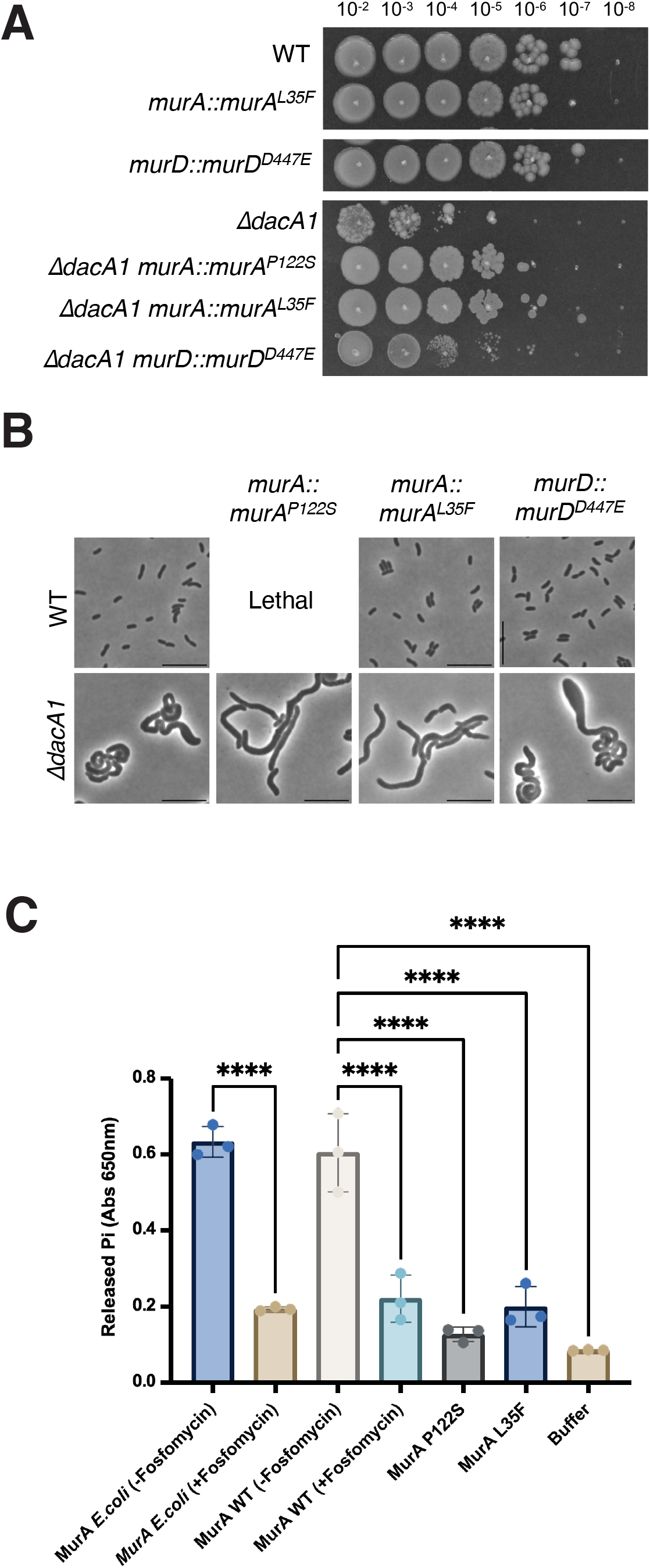
Hypomorphic mutants in the PG precursor synthesis pathway rescue *ΔdacA1* growth but not morphology. **(A)** Indicated strains are chromosomal replacements of *murA* and *murD* murA^P122S^, murA^L35F^ and murD^D447E^ respectively. Overnight cultures were plated on LB medium by spot-titering and incubated at 30°C overnight. **(B)** Overnight cultures of the indicated strains were subcultured 1:100 and grown at 37°C in LB medium for 1 hour, then backdiluted 1:100 in LB and incubated at 37°C for an additional 2 hours before imaging **(C)** The different MurA mutants were purified, and their activity was measured *in vitro* using a MurA assay kit which measures the inorganic phosphate released from the MurA enzymatic reaction. Statistical significance was determined by an ordinary one-way ANOVA, error bars represent the standard deviation of three replicates (raw data points also shown).

*ΔdacA1* cells with mutations *murA^P122S/L35F^* and *murD^D447E^* cells were 65%, 39% and 30% larger, and 92%, 72% and 9% longer, respectively. However, these MurA mutants exhibited ∼15% decrease in width compared to the Δ*dacA1* mutant, while the *murD^D447E^* mutant exhibited a 7% increase in width (Fig. 5B, Fig. S5). Thus, these mutations uncouple morphological aberrations from the ability to ultimately form a colony on a plate. Interestingly, mutations *murA^P122S^*and *murD^D447E^* made *ΔdacA1* sensitive to salt-free LB even though they rescued the Δ*dacA1* growth defect on LB medium, suggesting fine-tuned optimality in these mutations’ ability to rescue Δ*dacA1*. Further epistatic analysis using overexpression constructs revealed that *murC^A132T^* but neither *murA* nor *murD* mutations was dominant negative (Fig. S6). We hypothesize that this is due to MurC’s requirement for dimerization^22^ – the merodiploid expression of the mutated allele may result in MurC/MurC^A132T^ heterodimers with overall reduced function.

We next turned to an *in vitro* assay to biochemically probe *mur* mutant function. We thus purified MurA, MurA^P122S^ and MurA^L35F^ and measured their enzymatic activity using a MurA assay kit^23^. Both MurA mutants, MurA^P122S^ and MurA^L35F^ exhibited significantly lower activity than the WT, almost comparable to MurA treated with its inhibitor, fosfomycin. These results suggested that reducing cell wall synthesis alleviates Δ*dacA1* growth phenotypes. To further test this hypothesis, we chemically reduced precursor synthesis and measured growth of Δ*dacA1* cells on LB. Treatment of the Δ*dacA1* mutant with increasing concentrations of fosfomycin, which covalently binds to the active site of MurA, inhibiting its activity^24^, partially restored *ΔdacA1* cells growth on LB at concentrations between 5μg/mL and 15μg/mL (0.15 x - 0.45 x MIC) (Fig. S7). Thus, in a mechanistically unclear way, but likely tied to their reduced activity, mutations in MurA, MurC and MurD promote proper growth (colony formation), but not proper morphogenesis in Δ*dacA1*.

Depending on affinities of the multiple cell wall synthases for lipid II, reduced precursor synthesis may disproportionately affect the Rod system, divisome, and/or aPBPs. To test whether the observed fosfomycin rescue affect was due to selective reduction in the activity of one of these synthases, we treated WT and *ΔdacA1* cells with sub-MIC concentrations of rod complex inhibitors (A22, MP265 and mecillinam), the aPBP inhibitor moenomycin or the PBP3 inhibitor aztreonam. We found that independently inhibiting the different PG synthases did not result in *ΔdacA1* rescue (Fig. S8A). To next test if generalized PG synthesis inhibition resulted in Δ*dacA1* rescue, we treated WT and *ΔdacA1* cells with sub-MIC concentrations of A22 and Moenomycin simultaneously. Consistent with our prediction, preventing generalized PG synthesis partially rescued *ΔdacA1* in LB (Fig. S8B) in a very subtle and concentration-dependent, but reproducible way. Overall, these data additionally support a model where the dacA1 mutant suffers from an imbalance between cell wall synthesis and degradation, and can be rescued by either reducing PG synthesis (partial rescue), or enhancing PG degradation (full rescue).

### *Δ*dacA1 salt sensitivity is partially rescued by interfering with C55 metabolism and exacerbated by increased PG synthesis

Our data suggest that either reducing cell wall synthesis or increasing cell wall degradation can rescue growth of the *dacA1* mutant. Why then is the mutant salt-sensitive in the first place? In addition to a proton motive force for energy generation, *V. cholerae* also generates a sodium motive force to power its flagellum, and, as recently suggested, to potentially drive the essential recycling reaction of the undecaprenol pyrophosphate cell wall precursor carrier ^25^. Thus, growth in salt-free LB might limit *V. cholerae’s* capacity for optimal cell wall synthesis due to the absence of a sodium motive force; conversely, growth in LB might enhance PG synthesis. If this was the case, and enhanced PG synthesis in LB was the culprit for *ΔdacA1* defects, we would for example expect a mutant in the sodium-dependent undecaprenol pyrophosphate translocase, Vca0040, to suppress Δ*dacA1* phenotypes. To thus test whether there was a genetic relationship between *vca0040* and *dacA1*, we created a double mutant and tested its growth on LB. Consistent with our hypothesis, we observed that deleting *vca0040* in a *ΔdacA1* background partially restored growth (consistent 10-fold increase in plating efficiency), and complementation with an IPTG inducible copy of *vca0040* demonstrated it was in fact due to the absence of the translocase and not a polar effect (Fig. 6A, Fig. S9). These results suggest that VC_A0040 does contribute to the NaCl dependent toxicity in the ΔdacA1 mutant; however, it is not the only factor involved.

**Figure 6.**
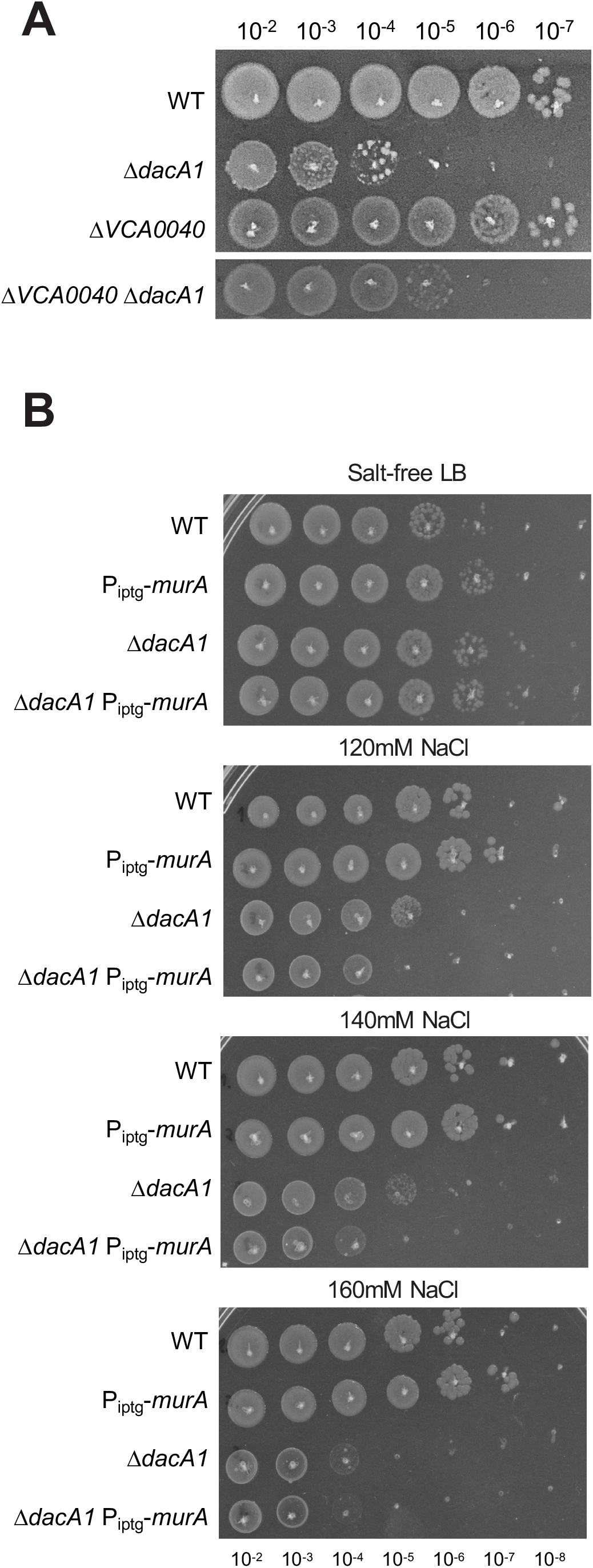
PG precursor recycling and synthesis modulate *dacA1* mutant fitness. **(A)** Overnight cultures of the indicated mutants (see main text for details) were plated on LB medium via spot-titering and incubated at 30°C overnight. **(B)** Overnight cultures of WT, *ΔdacA1* and both backgrounds harbouring an IPTG inducible copy of *murA* were plated on salt-free LB with increasing concentrations of NaCl (120mM, 140mM and 160mM) and incubated at 30°C overnight.

Next, we tested whether there is a relationship between NaCl toxicity and the availability of PG precursors. To this end, we overexpressed MurA (which results in upregulation of PG synthesis)^14^ in a *ΔdacA1* background and evaluated its effect of growth at different NaCl concentrations. We found that overexpression of MurA in *ΔdacA1* cells made them more susceptible to lower concentrations of NaCl (120mM) when compared to *ΔdacA1* cells not overexpressing MurA (which become susceptible to NaCl at a concentration of 140mM only) (Fig. 6B, Fig. S10). This again subtle, but reproducible phenotype indicates that increased availability of PG precursors, in addition to Na+ in the culture medium are implicated in the growth defects observed in *ΔdacA1* cells.

### The relationship between PBP5 and endopeptidases is conserved in *E. coli*

DacA1 homologues are collectively essential for proper cell shape maintenance in *E.coli*, and their absence leads to aberrant and irregular morphologies, similar to the *V. cholerae dacA1* mutant ^3, 26, 27^. To investigate if increased endopeptidase activity rescues the morphological defects caused by carboxypeptidase deficiency in *E. coli*, we overexpressed ShyA and ShyA^R115W^ in a strain lacking 4 DD-CPases, DacA, PbpG, DacB and DacD (Strain CS446-1^26^), hereafter referred to as Δ4. The Δ4 strain exhibits well-characterized, severe morphological defects, quantifiable as a drastic increase in cell area^28^. These defects could be almost fully suppressed by overexpressing ShyA (Fig. 7AB). We also attempted to rescue Δ4_EC_ by overexpressing our mutant ShyA^R115W^, however, this resulted in severe growth defects in both WT and *Δ*4 (Fig. S3). Thus, morphological defects associated with carboxypeptidase deficiency in *E. coli* can be rescued by increasing endopeptidase activity, indicating that the role of carboxypeptidases in maintaining the balance between cell wall synthesis and EP-mediated degradation is conserved.

**Figure 7.**
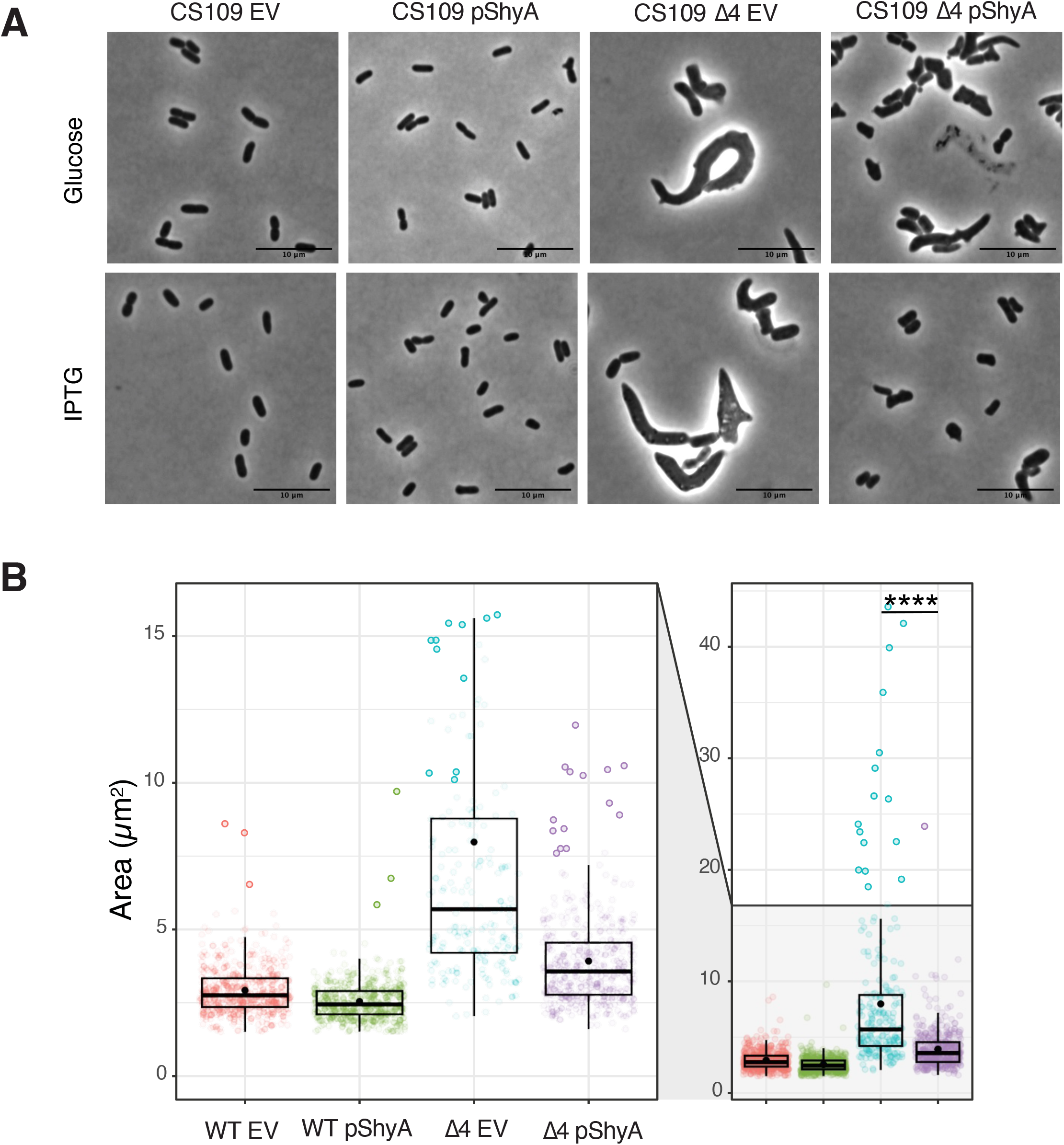
ShyA overexpression rescues morphological defects associated with carboxypeptidase insufficiency in *E. coli*. **(A)** *E. coli* strains CS109 and Δ4 (CS109 Δ*dacA ΔdacB ΔdacC ΔpbpG*) containing either an empty vector (pHL100) or a vector containing an IPTG-inducible copy or ShyA (pHL100-ShyA) were incubated overnight at 37°C in LB medium with 0.2% glucose, sub-cultured 1:100 in LB and incubated for 2 hours, then back-diluted 1:100 in LB and grown for an additional 2 hours. 1 mM or 0.2% glucose was added to the cultures, phase contrast images were taken 3 hours after induction. (B) Microscopy images were segmented with Omnipose and analyzed for area, length, and width with MicrobeJ. Statistical significance was assessed via Welch’s t-test. ****, P < 0.0001.

## Discussion

In this study, we have shown that defects associated with carboxypeptidase insufficiency can be alleviated by either enhancing cell wall degradation, or by reducing cell wall synthesis. Fundamentally, this suggests that a key role of carboxypeptidases is to ensure proper balance between PG synthesis and degradation (Fig. 8). Our data add support to a model that increased cell wall synthesis can be detrimental if not properly balanced by degradation. Studies in *Bacillus subtillis* described a similar phenomenon where accumulation of PG precursors disturbed the balance between PG synthesis and degradation which could be restored by increased D,L endopeptidase activity (LytE) or in its absence by mutations that downregulated PBP1^29^. It is important to note, however, that enhanced degradation seems to be the superior way to rescue Δ*dacA1* defects – while enhancing EP activity completely suppressed Δ*dacA1* growth and morphology, reduction of synthesis either chemically or through *mur* hypomorphs only partially rescued growth but did not restore morphology. This may either indicate that more severe trade-offs are associated with downregulating synthesis than with upregulating degradation, or that ultimately, crosslink cleavage is required and reducing synthesis indirectly allows effective cleavage of fewer crosslinks via other EPs (the *V. cholerae* genome encodes 9 such enzymes) with lower activity.

**Figure 8.**
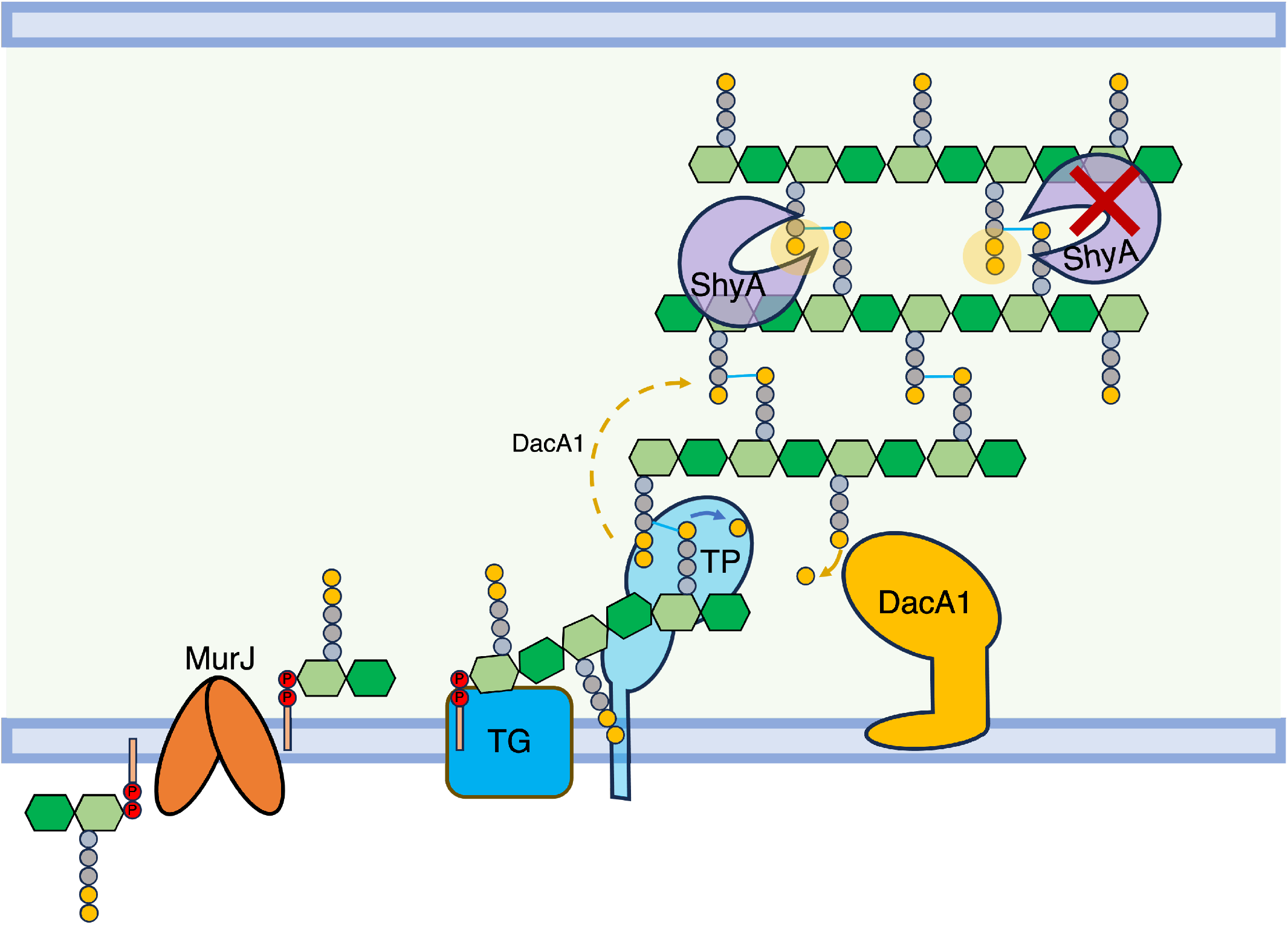
Model of DacA1’s role in balancing PG synthesis and degradation. DacA1 removes the 5^th^ D-ala residue of the PG side stem and determines which side stem can be used as donor for transpeptidation. By removing the 5^th^ D-ala residue, DacA1 also allows ShyA to cleave crosslinks between strands by producing its preferred substrate, tetrapeptide-containing side stems. In the absence of DacA1, there is an increased amount of pentapeptides in the PG sacculus, hindering ShyA’s ability to cleave crosslinks. This reduction in endopeptidase activity disrupts the balance between PG synthesis and degradation.

Carboxypeptidase function has been linked to cell shape maintenance in *E. coli* based on the observation that their absence results in severe morphological defects^3, 27, 28^. This has been attributed to excessive transpeptidation and incorrect crosslinking as a consequence of an increase in pentapeptide side stems, which can serve as both donors and acceptors in transpeptidation (while the majority of PG crosslinks consist of tetrapeptide and can only serve as donor). However, pentapeptide *per se* does not seem to be detrimental (the *V. cholerae* Δ*dacA1* mutant exhibits WT growth and morphology in salt-free LB, despite increased pentapeptide content), suggesting the presence of alleviating factors, and perhaps suggesting another role for carboxypeptidases^27^. Our data here suggest that instead of (or perhaps in addition to) promoting incorrect crosslinking, pentapeptide might impede EP-mediated PG cleavage (by increasing the amount of the pentapeptide substrate that is disfavoured by ShyA^6^) (Fig. S11), resulting in reduced insertion of nascent PG. Since *ΔdacA1* phenotypes do not mimic simple PG synthase inhibition, however, it is likely that the lack of cleavage has more complex architectural consequences. For example, we and others have previously shown that at least in *V. cholerae* and *B. subtilis* (unlike *E. coli*^7^), EP activity is not absolutely required for PG synthesis, but rather for the directional post-synthesis insertion of PG during cell elongation. Though highly speculative, this may point towards Holtje’s “make before break” model^30^, where PG is constructed as a second layer beneath the first and then inserted by the action of EPs. If this scenario is correct, locally reduced EP activity might create expansive stretches of PG with pentapeptide-rich second layers, locally restricting cell elongation, which could result in the aberrant shapes that are typically associated with lack of carboxypeptidases – like an air balloon being inflated with patches of adhesive tape on its side. This model is consistent with studies done in *E. coli*, where uncharacterized mutations associated with the endopeptidases AmpH and MepA suppress the severe morphological defects in strains missing 3 or 4 PBPs (including DacA)^31^.

Our data also reveal a potential reason for the salt sensitivity of the Δ*dacA1* mutant in *V. cholerae*.

In contrast to this mutant’s phenotype, cell wall mutants are often sensitive to low salt conditions due to their inability to withstand their internal osmotic pressure ^32^. Our data suggest that surplus PG synthesis is partially responsible for the *dacA1* defects, and perhaps absence of NaCl reduces PG synthesis. *V. cholerae* can generate a sodium motive force (SMF) via its Na(+)-NQR enzyme^33^. Recent work has proposed that SMF is important for cell wall precursor recycling as it can power the C55-P translocase encoded in gene *Δvca0040*^25^, and deletion of *vca0040* indeed reduced salt-sensitivity of the *dacA1* mutant, albeit very modestly. It is formally possible that other cell wall functions across the inner membrane, such as Lipid II flipping by MurJ may also rely on the SMF, boosting PG synthesis in the presence of surplus Na^+^. MurJ is known to require membrane potential, but not necessarily an H+ gradient to function. The specific ions involved in lipid II transport by MurJ depend on the organism; for instance, *Thermosipho africanus* requires chloride ions for MurJ function, while MurJ from *E. coli* uses the proton motive force^34^. Interestingly, a recent report demonstrated sensitivity to very high salt (750 mM) in the *dacA* mutant in *E. coli* as well^35^. Since we do not expect a sodium motive force to promote cell wall synthesis in *E. coli,* this sensitivity may rather reflect non-specific physiology, e.g., disruption of protein-protein interactions that are detrimental only in the absence of PBP5.

More broadly, or data imply that the balance between PG synthesis and degradation does not necessarily have to be maintained by direct protein-protein interactions (tight complexes between hydrolases and synthases), since the same end-result can be achieved by upregulating degradation in diverse ways and downregulating synthesis in diverse ways. A physical uncoupling between PG synthesis and degradation may also explain the high redundancy in synthases and hydrolases, as this allows for the maintenance of this balance under varying environmental conditions without relying on strong and specific protein complexes.

## Acknowledgements

This work was supported by NIH grant R01GM130971 (TD). We thank Matt Jorgenson and Kevin Young (University of Arkansas) for the gift of CS109 and the Δ4 derivative.

## Methods

### Bacterial strains, media, and growth conditions

All *V. cholerae* strains used in this study are derivatives of El Tor strains N16961 and C6706 (Table S1). *V. cholerae* cultures were grown by shaking (200rpm) at 30°C in LB broth (Tryptone 10g/L, Yeast extract 5g/L, Sodium Chloride 10g/L) or salt-free LB (Tryptone 10g/L, Yeast extract 5g/L) unless otherwise indicated. 200µg/mL of streptomycin was also added to the cultures (the *V. cholerae* strains used are streptomycin resistant). When appropriate, the overnight cultures were prepared with 0.2% glucose to prevent the leaky expression from the P_IPTG_ promoter. For growth curve experiments, overnight cultures were diluted 100-fold into 1 mL of the appropriate growth medium containing streptomycin and incubated at 37°C with shaking in 100-well honeycomb wells in a Bioscreen plate reader (Growth Curves America) which recorded optical density at 600 nm (OD600) in 10-minute intervals. *E. coli* cultures were grown by shaking (200rpm) at 37°C in LB broth unless otherwise stated. When appropriate, antibiotics were used in the following concentrations: carbenicillin (100 µg/mL), kanamycin (50 µg/mL) and chloramphenicol (20 µg/mL). Genes under P_IPTG_ regulation were induced with 1mM IPTG and repressed with 0.2% glucose in both *V. cholerae* and *E. coli*.

### Cloning, vectors, and strain construction

A summary of the plasmids and primers used in this study can be found in Supplementary Table 2. *E. coli* DH5α *λ pir* was used for general cloning, *E. coli* SM10 and MFD *λ pir* were used for conjugation into *V. cholerae*. Plasmids were built using isothermal assembly ^36^ and verified by Sanger Sequencing.

Chromosomal in-frame deletions were generated using the allelic exchange vector pTOX5 cmR/msqR^37^. Regions flanking the gene to be deleted (500-700bp) were amplified from *V. cholerae* N16961 genomic DNA, cloned into the suicide vector pTOX5 and transformed into *E. coli* DH5α. Chloramphenicol resistant colonies were verified by colony PCR using primers MA83 and MA84 and verified via Sanger Sequencing. Correct constructs were selected and transformed into *E. coli* SM10 (Donor) and Conjugated into *V. cholerae* (recipient) by spotting 15µL of the Donor and 15µL of the recipient onto a LB plate with 1% Glucose and incubating for 4h at 37°C. Transconjugants were selected on LB plates with chloramphenicol (20 µg/mL), streptomycin (200 µg/mL) and 1% Glucose after incubation at 30°C overnight. Chloramphenicol-and streptomycin-resistant colonies were grown without selective pressure in microcentrifuge tubes with 1mL LB broth with 1% Glucose and incubated for 3h at 37°C. 2-3 candidates were then streaked on M9 minimal medium supplemented with 0.2% (wt/vol) Casamino Acids, 0.5 mM MgSO4, 0.1 mM CaCl2, 25 µM iron chloride in 50 µM citric acid and 2% rhamnose, and grown at 30°C. Single colonies were further purified by streaking onto M9-Rhamnose plates and deletions were verified by PCR using flanking and internal primers for each gene and validated by whole genome sequencing. Given *ΔdacA1* salt-sensitivity, all cloning steps with LB media were replaced with salt-free LB.

For complementation of deletion strains, genes were amplified from *V. cholerae* N16961 with primers designed to introduce a strong ribosome-binding site (RBS) (AGGAGG) and cloned into pTD101 a pJL1^38^ derivative, which contains *lacI^q^*, a multiple cloning site downstream of an IPTG (isopropyl-β-D-thiogalactopyranoside)-inducible promoter (P_IPTG_) and allows chromosomal insertions of the gene of interest into the native *lacZ locus* in *V. cholerae*. Successful insertions were verified by PCR using lacZ flaking primers (MA106-107).

To replace the chromosomal copies of *murA* and *murD* with the mutants isolated from the suppressor screen, we used the allelic exchange vector pTOX5 as described above, however, instead of just using the *murA* and *murD* flanking regions, we used the gDNA from the suppressor mutants and used primers that annealed 500bp upstream and 500bp downstream to amplify the mutant gene with its corresponding flanking regions for insertion into the chromosomal WT locus. Successful integration was validated by PCR amplification and Sanger sequencing.

ShyA^R115W^ mutant constructs were generated by amplifying genomic DNA from the suppressor strain and cloned into pTD101 and pHLmob100. Constructs were verified using Sanger Sequencing.

### TnSeq

TnSeq was conducted as described before^39, 40^; Briefly, cultures of strains MA1 and MA73 were mated with donor strain *E.coli* MFD λ pir, which contains the pSC189 suicide plasmid encoding the mariner transposon. Approximately 200,000 colonies/replicate were recovered from plates containing Kan50 to select for transposon insertions and 1mM IPTG to allow the expression of ShyA^L109K^. Libraries were then resuspended in 30 mL of LB broth, and 1/5 of the culture was used for genomic DNA (gDNA) extraction and the rest was frozen in 30% glycerol at −80°C. Samples were prepared for sequencing as follows. The extracted gDNA was sheared by sonication (9 seconds, 30% amplitude), followed by blunting (Blunting Enzyme Mix, NEB), A-tailing, ligation of specific adaptors and PCR amplification of the transposon-DNA junctions using transposon-and adaptor-specific primers. Libraries were sequenced using Illumina MiSeq as described previously ^41^. To determine gene essentiality, data analysis was performed using the Matlab-based pipeline ARTIST^40^. Genetic regions predicted to be conditionally essential/enriched were inspected using the genome browser Artemis^42^ and insertion plots were generated using the tidyverse package in R.

### PG purification and Remazol blue assay of hydrolase activity

Peptidoglycan sacculi were purified as described previously^43^. Briefly, 1 L of WT and *ΔdacA1* cultures were harvested and resuspended in boiled 5% SDS for 30 minutes. The cell lysate was pelleted by ultracentrifugation at 130,000g for 60 minutes at room temperature and resuspended in 30mL of MilliQ water at 50°C, the pellets were washed 4 times before being resuspended in 100mM Tris HCl pH 7.5 with 100 ug of trypsin and CaCl2 to a final concentration of 10 mM and incubated at 37°C overnight. On the next day, samples were boiled for 15 min to inactivate the Trypsin, washed twice with 30 mL of warm MilliQ water and centrifuged at 130,000g for 30 minutes at 20°C. The pellet was then resuspended in 500uL of MilliQ water and stored at −80°C.

Purified sacculi were stained with Remazol Brilliant Blue (RBB) as described previously^44^. In a 15mL conical tube, 1mL of sacculi was mixed with 0.4 mL of 0.2M Remazol-brilliant tube (Sigma, R8001), 0.3 mL of 5M NaOH and 4.3 mL of MilliQ water and incubated at 37°C overnight on a rocker table. To neutralize the pH and wash away the residual dye, 0.4 mL of 5M HCl was added alongside 0.75 mL of 10X PBS, samples were then ultracentrifuged at 400,000g for 20 min at 22°C and resuspended in MilliQ water, the samples were washed multiple times until the supernatant ran clear. To eliminate possible contamination with lysozyme, the stained sacculi were incubated at 65°C for 3 hours and stored in 10%glycerol at −20°C.

Endopeptidase activity reactions were prepared with 200µL of Pulldown buffer^5^ (20 mM Tris/maleate (pH 6.8), 30 mM Nacl, 10 mM MgCl, 1 mM DTT, 0.1% Triton X-100), 50uL of RBB-stained sacculi and 10 µg of lysozyme purified ShyA and ShyA^L109K^, and incubated at 37°C for 3h. The reactions were stopped by adding 1% SDS and washed by ultracentrifugation at 80,000g for 15 min. Supernatants were removed and A_585_ measured in 96-well plates using a microplate reader.

### Microscopy and image analysis

Cells were grown as previously described and imaged under phase contrast on an agarose pad (0.8% agarose in LB or salt-free LB) using a Leica DMi8 inverted microscope. Images were segmented using Omnipose^45^ using the bact_phase_omni model, masks were exported as .png and imported into MicrobeJ^46^ for image analysis and quantification using the default parameters. Significant differences were determined using Welch’s two-sample t-test. Plots were made using the R packages ggplot2 and tidyverse.

### Spontaneous suppressor identification

*ΔdacA1* cells were grown overnight in SF media, 1mL of overnight culture was spread on a BioAssay Square Dish (Thermo Scientific™ Nunc™) containing 200 mL of LB medium with Streptomycin 200 µg/mL. This experiment was performed 3 independent times, for each instance, 16 colonies were selected from the plate, to validate the stability of the mutations each colony was grown overnight in salt-free LB and then in LB agar. Genomic DNA from was extracted and samples were sent to SeqCenter (Pittsburg, PA) for whole genome sequencing and variant calling analysis.

### Protein purification

MurA, MurA^P122S^ and MurA^L35F^ genes were amplified from N16961 gDNA and the gDNA from the mutants obtained from the suppressor screen, and cloned into pET28a downstream of 6xHis-SUMO Tag. Plasmids were verified by PCR and Whole Plasmid Sequencing through Plasmidsaurus (Eugene, OR). *E. coli* BL21 (DE3) (Novagen) was transformed with the resulting recombinant plasmids (pET28a-MurA, pET28-MurA^P122S^ and pET28a-MurA^L35F^). Overnight cultures (10 mL) were used to inoculate 1L of LB with Kanamycin (50 µg/mL) and incubated at 37°C with vigorous shaking (220 RPM) until they reached an OD_600_ between 0.6 and 0.8. Cultures were induced with 1mM IPTG at 18°C and 180 RPM overnight. Harvested cells were pelleted and resuspended in 30 mL of cold Purification Buffer (20mM Tris pH 7.5, 150 mM NaCl, 1mM PMSF), and lysed by sonication. Lysates were cleared by centrifugation at 31,000g for 4 minutes at 4°C, and loaded onto a HisPur Cobalt column (Thermo Scientific; Catalog No. 89964) and washed multiple times with Purification Buffer until protein was undetectable by the Bradford reagent. The bead slurry was then transferred to a 5 mL microtube with 60 µL of ULP1 Sumo protease and digested overnight at 4°C rotating. Protein was eluted the next day with 20 mL of Purification Buffer. Samples were analysed by SDS-page with Coomassie Blue Stain and then Concentrated with a 30KD Amicon concentrator (Millipore) to 5 mL. Concentrated samples were then passed through a HiLoad 16/600 Superdex 75pg gravity column using and ÄKTA pure^TM^ chromatography system (Cytiva). ShyA and ShyA^L109K^, were purified as previously described^12^.

### MurA enzymatic activity

The enzymatic activity of MurA, MurA^P122S^ and MurA^L35F^, was measured using a bacterial MurA Assay (Profoldin, Hudson MA) based on the spectrophotometric measurement of inorganic phosphate released from the enzymatic reaction catalysed by MurA. This reaction transfers enolpyrupate from phosphoenolpyruvate (PEP) to uridine diphospho-N-acetylglucosamine (UNAG), which results in the generation of enolpyruvyl-UDPN-acetylglucodamine (EP-UNAG) and inorganic phosphate (Fig. 5).

For this experiment MurA*^E.coli^*provided by the manufacturer was used as a control, and 1µL of 50mg/mL Fosfomycin was added to the WT versions of MurA to validate the ability of the test to detect reduced activity of MurA. The reactions were performed by triplicate following the manufacturer’s protocol. Briefly 5000mM of each protein was resuspended in the reaction buffer (for 1 reaction: 6.6 µL of 10X buffer and 52.2 µL of H_2_O) and the enzyme substrate was prepared (for 1 reaction: 0.66 µL of 100X PEP, 0.66 µL of 100X UGN and 5.28 µL of H_2_O), the reactions were prepared by adding 54 µL of resuspended MurA and 6 µL of the enzyme substrate to a 96-well plates, samples were incubated at 37 °C for 60 min. For inorganic phosphate detection, 90µL of Dye MPA3000 was added and samples were incubated for 5 mins before measuring light absorbance at 650nm in a 96-well plate reader.

## Data availability

This study includes no data deposited in external repositories.

## Bibliography

1. Cho, H. et al. Bacterial cell wall biogenesis is mediated by SEDS and PBP polymerase families functioning semi-autonomously. Nat. Microbiol. 1, 1–8 (2016).

2. Mueller, E. A. & Levin, P. A. Bacterial Cell Wall Quality Control during Environmental Stress. mBio 11, (2020).

3. Nelson, D. E. & Young, K. D. Penicillin Binding Protein 5 Affects Cell Diameter, Contour, and Morphology of Escherichia coli. J. Bacteriol. 182, 1714–1721 (2000).

4. Ghosh, A. S., Chowdhury, C. & Nelson, D. E. Physiological functions of D-alanine carboxypeptidases in Escherichia coli. Trends Microbiol. 16, 309–317 (2008).

5. Möll, A. et al. A D, D-carboxypeptidase is required for Vibrio cholerae halotolerance. Environ. Microbiol. 17, 527–540 (2015).

6. Dörr, T., Cava, F., Lam, H., Davis, B. M. & Waldor, M. K. Substrate specificity of an elongation-specific peptidoglycan endopeptidase and its implications for cell wall architecture and growth of Vibrio cholerae. Mol. Microbiol. 89, 949–962 (2013).

7. Singh, S. K., SaiSree, L., Amrutha, R. N. & Reddy, M. Three redundant murein endopeptidases catalyse an essential cleavage step in peptidoglycan synthesis of Escherichia coli K12. Mol. Microbiol. 86, 1036–1051 (2012).

8. Hashimoto, M., Ooiwa, S. & Sekiguchi, J. Synthetic Lethality of the lytE cwlO Genotype in Bacillus subtilis Is Caused by Lack of d,l-Endopeptidase Activity at the Lateral Cell Wall. J. Bacteriol. 194, 796–803 (2012).

9. Murphy, S. G. et al. Endopeptidase Regulation as a Novel Function of the Zur-Dependent Zinc Starvation Response. mBio 10, (2019).

10. Singh, S. K., Parveen, S., SaiSree, L. & Reddy, M. Regulated proteolysis of a cross-link– specific peptidoglycan hydrolase contributes to bacterial morphogenesis. Proc. Natl. Acad. Sci. 112, 10956–10961 (2015).

11. Shin, J.-H. et al. Structural basis of peptidoglycan endopeptidase regulation. Proc. Natl. Acad. Sci. U. S. A. 117, 11692–11702 (2020).

12. Shin, J.-H., et al. Structural basis of peptidoglycan endopeptidase regulation. bioRxiv 843615 (2019) doi:10.1101/843615.

13. Dörr, T. et al. A Novel Peptidoglycan Binding Protein Crucial for PBP1A-Mediated Cell Wall Biogenesis in Vibrio cholerae. PLoS Genet. 10, e1004433 (2014).

14. Hernández, S. B., Dörr, T., Waldor, M. K. & Cava, F. Modulation of Peptidoglycan Synthesis by Recycled Cell Wall Tetrapeptides. Cell Rep. 31, 107578 (2020).

15. Rai, A. K. et al. Differentiation of gram-negative intermembrane phospholipid transporter function by fatty acid saturation preference. 2023.06.21.545913 Preprint at https://doi.org/10.1101/2023.06.21.545913 (2023).

16. Grimm, J. et al. The inner membrane protein YhdP modulates the rate of anterograde phospholipid flow in Escherichia coli. Proc. Natl. Acad. Sci. U. S. A. 117, 26907–26914 (2020).

17. Skarzynski, T. et al. Structure of UDP-N-acetylglucosamine enolpyruvyl transferase, an enzyme essential for the synthesis of bacterial peptidoglycan, complexed with substrate UDP-N-acetylglucosamine and the drug fosfomycin. Struct. Lond. Engl. 1993 4, 1465–1474 (1996).

18. de Oliveira, M. V. D., Furtado, R. M., da Costa, K. S., Vakal, S. & Lima, A. H. Advances in UDP-N-Acetylglucosamine Enolpyruvyl Transferase (MurA) Covalent Inhibition. Front. Mol. Biosci. 9, (2022).

19. Yu, H., Zhao, Y., Guo, C., Gan, Y. & Huang, H. The role of proline substitutions within flexible regions on thermostability of luciferase. Biochim. Biophys. Acta BBA - Proteins Proteomics 1854, 65–72 (2015).

20. Wamp, S. et al. MurA escape mutations uncouple peptidoglycan biosynthesis from PrkA signaling. PLOS Pathog. 18, e1010406 (2022).

21. Hummels, K. R. et al. Coordination of bacterial cell wall and outer membrane biosynthesis. Nature 615, 300–304 (2023).

22. Jin, H. et al. Structural Studies of Escherichia coli UDP-N-Acetylmuramate:l-Alanine Ligase. Biochemistry 35, 1423–1431 (1996).

23. Kurnia, D., Apriyanti, E., Soraya, C. & Satari, M. H. Antibacterial Flavonoids Against Oral Bacteria of Enterococcus Faecalis ATCC 29212 from Sarang Semut (Myrmecodia pendans) and Its Inhibitor Activity Against Enzyme MurA. Curr. Drug Discov. Technol. 16, 290–296 (2019).

24. Falagas, M. E., Vouloumanou, E. K., Samonis, G. & Vardakas, K. Z. Fosfomycin. Clin Microbiol. Rev. 29, 321–347 (2016).

25. Sit, B. et al. Undecaprenyl phosphate translocases confer conditional microbial fitness. Nature 613, 721–728 (2023).

26. Denome, S. A. Escherichia coli Mutants Lacking All Possible Combinations of Eight Penicillin Binding Proteins: Viability, Characteristics, and Implications for Peptidoglycan Synthesis. https://journals.asm.org/doi/epdf/10.1128/JB.181.13.3981-3993.1999?src=getftr doi:10.1128/JB.181.13.3981-3993.1999.

27. Nelson, D. E. & Young, K. D. Contributions of PBP 5 and dd-Carboxypeptidase Penicillin Binding Proteins to Maintenance of Cell Shape in Escherichia coli. J. Bacteriol. 183, 3055– 3064 (2001).

28. Peters, K. et al. The Redundancy of Peptidoglycan Carboxypeptidases Ensures Robust Cell Shape Maintenance in Escherichia coli. mBio 7, e00819–16 (2016).

29. Sassine, J., Sousa, J., Lalk, M., Daniel, R. A. & Vollmer, W. Cell morphology maintenance in Bacillus subtilis through balanced peptidoglycan synthesis and hydrolysis. Sci. Rep. 10, 17910 (2020).

30. Höltje, J. V. Growth of the stress-bearing and shape-maintaining murein sacculus of Escherichia coli. Microbiol. Mol. Biol. Rev. MMBR 62, 181–203 (1998).

31. Laubacher, M. E., Melquist, A. L., Chandramohan, L. & Young, K. D. Cell Sorting Enriches Escherichia coli Mutants That Rely on Peptidoglycan Endopeptidases To Suppress Highly Aberrant Morphologies. J. Bacteriol. 195, 855–866 (2013).

32. Weaver, A. I. et al. Lytic transglycosylases mitigate periplasmic crowding by degrading soluble cell wall turnover products. eLife 11, e73178 (2022).

33. Häse, C. C. & Barquera, B. Role of sodium bioenergetics in Vibrio cholerae. Biochim. Biophys. Acta 1505, 169–178 (2001).

34. Kumar, S., Mollo, A., Rubino, F. A., Kahne, D. & Ruiz, N. Chloride Ions Are Required for Thermosipho africanus MurJ Function. mBio 14, e00089–23 (2023).

35. Choi, U., Park, S. H., Lee, H. B., Son, J. E. & Lee, C.-R. Coordinated and Distinct Roles of Peptidoglycan Carboxypeptidases DacC and DacA in Cell Growth and Shape Maintenance under Stress Conditions. Microbiol. Spectr. 0, e00014–23 (2023).

36. Gibson, D. G. et al. Enzymatic assembly of DNA molecules up to several hundred kilobases. Nat. Methods 6, nmeth.1318 (2009).

37. Lazarus, J. E. et al. A New Suite of Allelic-Exchange Vectors for the Scarless Modification of Proteobacterial Genomes. Appl. Environ. Microbiol. 85, (2019).

38. Miyata, S. T., Unterweger, D., Rudko, S. P. & Pukatzki, S. Dual Expression Profile of Type VI Secretion System Immunity Genes Protects Pandemic Vibrio cholerae. PLOS Pathog. 9, e1003752 (2013).

39. Yamaichi, Y. & Dörr, T. Transposon Insertion Site Sequencing for Synthetic Lethal Screening. in The Bacterial Nucleoid: Methods and Protocols (ed. Espéli, O.) 39–49 (Springer, 2017). doi:10.1007/978-1-4939-7098-8_4.

40. Pritchard, J. R. et al. ARTIST: High-Resolution Genome-Wide Assessment of Fitness Using Transposon-Insertion Sequencing. PLoS Genet. 10, e1004782 (2014).

41. Ferrières, L. et al. Silent mischief: bacteriophage Mu insertions contaminate products of Escherichia coli random mutagenesis performed using suicidal transposon delivery plasmids mobilized by broad-host-range RP4 conjugative machinery. J. Bacteriol. 192, 6418–6427 (2010).

42. Carver, T., Harris, S. R., Berriman, M., Parkhill, J. & McQuillan, J. A. Artemis: an integrated platform for visualization and analysis of high-throughput sequence-based experimental data. Bioinformatics 28, 464–469 (2012).

43. Alvarez, L., Hernandez, S. B., de Pedro, M. A. & Cava, F. Ultra-Sensitive, High-Resolution Liquid Chromatography Methods for the High-Throughput Quantitative Analysis of Bacterial Cell Wall Chemistry and Structure. in Bacterial Cell Wall Homeostasis: Methods and Protocols (ed. Hong, H.-J.) 11–27 (Springer, 2016). doi:10.1007/978-1-4939-3676-2_2.

44. Zhou, R., Chen, S. & Recsei, P. A dye release assay for determination of lysostaphin activity. Anal. Biochem. 171, 141–144 (1988).

45. Cutler, K. J. et al. Omnipose: a high-precision morphology-independent solution for bacterial cell segmentation. Nat. Methods 19, 1438–1448 (2022).

46. Ducret, A., Quardokus, E. M. & Brun, Y. V. MicrobeJ, a tool for high throughput bacterial cell detection and quantitative analysis. Nat. Microbiol. 1, 1–7 (2016).

47. Schaub, R. E. & Dillard, J. P. Digestion of Peptidoglycan and Analysis of Soluble Fragments. Bio-Protoc. 7, e2438 (2017).

